# Early Inflammation and Interferon Signaling Direct Enhanced Intestinal Crypt Regeneration after Proton FLASH Radiotherapy

**DOI:** 10.1101/2024.08.16.608284

**Authors:** Tristan L. Lim, Clara Morral, Ioannis I. Verginadis, Kyle Kim, Leo Luo, Caitlin J. Foley, Michele M. Kim, Ning Li, Benjamin Yoshor, Kizito Njah, Mary Putt, Seyyedeh Azar Oliaei Motlagh, Anastasia Velalopoulou, Priyanka Chowdhury, Sandra Bicher, Denisa Goia, Christopher J. Lengner, Jeffery L. Wrana, Constantinos Koumenis, Andy J. Minn

**Affiliations:** Department of Radiation Oncology, Perelman School of Medicine, University of Pennsylvania, Philadelphia, PA, USA; Department of Biomedical Sciences, University of Pennsylvania School of Veterinary Medicine, University of Pennsylvania, Philadelphia, PA, USA; Centre for Systems Biology, Lunenfeld-Tanenbaum Research Institute, Mount Sinai Hospital, Toronto, Ontario, Canada; Department of Biostatistics, Epidemiology & Informatics, Perelman School of Medicine, University of Pennsylvania, Philadelphia, PA, USA; Department of Molecular Genetics, University of Toronto, Toronto, Ontario, Canada; Abramson Family Cancer Research Institute, Perelman School of Medicine, University of Pennsylvania, Philadelphia, PA, USA; Mark Foundation Center for Immunotherapy, Immune Signaling, and Radiation, University of Pennsylvania, Philadelphia, PA, USA

**Author notes:** Contributed equally to this work (Tristan L. Lim, Clara Morral, Ioannis I. Verginadis).

## Abstract

Ultra-high dose rate (“FLASH”) radiotherapy (≥40-60 Gy/s) is a promising new radiation modality currently in human clinical trials. Previous studies showed that FLASH proton radiotherapy (FR) improves toxicity of normal tissues compared to standard proton radiotherapy (SR) without compromising anti-tumor effects. Understanding this normal tissue sparing effect may offer insight into how toxicities from cancer therapy can be improved. Here, we show that compared to SR, FR resulted in improved acute weight recovery and survival in mice after whole-abdomen irradiation. Improved morbidity and mortality after FR were associated with greater proliferation of damage-induced epithelial progenitor cells followed by improved tissue regeneration. FR led to the accelerated differentiation of revival stem cells (revSCs), a rare damage-induced stem cell required for intestinal regeneration, and to qualitative and quantitative changes in activity of signaling pathways important for revSC differentiation and epithelial regeneration. Specifically, FR resulted in greater infiltration of macrophages producing TGF-β, a cytokine important for revSC induction, that was coupled to augmented TGF-β signaling in revSCs. In pericryptal fibroblasts, FR resulted in greater type I IFN (IFN-I) signaling, which directly stimulates production of FGF growth factors supporting revSC proliferation. Accordingly, the ability of FR to improve epithelial regeneration and morbidity was dependent on IFN-I signaling. In the context of SR, however, IFN-I had a detrimental effect and promoted toxicity. Thus, a tissue-level signaling network coordinated by differences in IFN-I signaling and involving stromal cells, immune cells, and revSCs underlies the ability of FLASH to improve normal tissue toxicity without compromising anti-tumor efficacy.

**One Sentence Summary:** FLASH radiation improves normal tissue toxicity without compromising anti-tumor efficacy through a stroma, immune, and revSC signaling network coordinated by IFN-I.

## INTRODUCTION

Reducing treatment-related morbidity and mortality remains a significant challenge within the field of cancer therapeutics (*1–4*). For example, radiotherapy (RT), which is utilized during treatment of more than half of all cancer patients, is associated with significant toxicities that constrain dose escalation to can improve efficacy and/or impact the quality of life for long-term survivors. However, several preclinical studies in rodents and larger animals have demonstrated that increasing the dose rate of RT to levels that far exceed standard dose rates can unexpectedly decrease toxicity to healthy tissue without appreciably compromising antitumor effects (*5–9*). This form of ultrahigh dose-rate radiation (≥40-60Gy/s) is called “FLASH” radiation and can deliver a therapeutic dose in a single treatment setting or hypofractionated setting. This is a radical departure from conventional (or standard) dose-rate radiation (≤0.03Gy/s) that requires splitting the same dose over a course of many treatments lasting several weeks. Splitting the dose (fractionation) is often needed in order to minimize toxicity but can also compromise the effectiveness of tumor control (*10*).

The ability of FLASH to deliver an entire course of radiation in a single dose that decreases normal tissue toxicity yet preserves anti-tumor efficacy is called the “FLASH effect.” Accordingly, FLASH is poised to profoundly change how RT is delivered to cancer patients, a goal that has prompted several clinical trials (*11, 12*). However, the underlying biology of the FLASH effect is largely unknown (*13*). Several studies have hypothesized that differential oxygen depletion when irradiating at ultrahigh dose-rates may underlie the selective advantage to healthy tissues with FLASH (*14–16*). Additionally, FLASH may decrease radiation exposure to circulating immune cells, thereby reducing inflammatory damage and achieving better tumor control and/or post-injury tissue repair (*17, 18*). Nonetheless, targeted investigations of these pathways have not yielded sufficient evidence to individually explain the FLASH effect, suggesting a complex interplay of several biological mechanisms (*19–23*). Understanding key mechanisms underlying the FLASH effect is likely critical in the clinical development of this promising new radiation modality with broad implications for improving cancer therapies in general (*24*).

The small intestine is a radiosensitive organ that offers a well-defined cellular architecture to study post-injury tissue regeneration (*25*). Injury to the intestinal epithelium is repaired, and barrier and absorptive functions are restored, through a hierarchical tissue structure that involves resident intestinal stem cell (ISC) niches in the epithelial crypts, as well as rare populations of damage-induced progenitor cells known as revival stem cells (revSCs) that are marked by the gene *Clu* (*26*). After intestinal damage from radiation, revSCs are thought to emerge from quiescence into a proliferative state to promote epithelial differentiation, while at the same time replenishing the homeostatic stem cell pool identified by *Lgr5* and *Olfm4*. This rapid induction and proliferation of *Clu*+ revSCs is thought to require p53 and coordination of multiple signaling pathways instigated by various stromal and immune cells within the lamina propria (*27*). This includes induction of revSCs by TGF-β produced by monocytes and macrophages, followed by proliferation of revSCs supported by FGF and EGF family of growth factors secreted by fibroblasts (*28, 29*). Recent evidence also demonstrates a role for type I interferon (IFN-I) in balancing differentiation and maintenance of ISCs. While IFN-I signaling can exert positive effects favoring epithelial regeneration, IFN-I signaling resulting from pathology or infection can also lead to attrition of the ISC pool in favor of differentiation (*30–33*).

In this study, we investigated the effects of single-dose mouse whole-abdomen FLASH and standard proton RT (FR and SR, respectively) on intestinal regeneration and associated morbidity and mortality. We show that single-dose whole-abdomen FR improves morbidity and mortality compared to SR. This enhanced recovery correlated with accelerated crypt regeneration linked to a network of key signaling pathways between stromal cells, immune cells, and revSCs. IFN-I signaling plays a key role specifically in the FLASH effect by coordinating this network, regulating ISC proliferation and revSC differentiation, and improving morbidity. Conversely, IFN-I signaling generated after SR has the opposite effect and promotes increased tissue toxicity.

## RESULTS

### Improved weight recovery and survival after FLASH is associated with accelerated crypt regeneration

We previously compared how FR versus SR might differentially impact epithelial regeneration after whole-abdomen irradiation of female C57BL/6 wildtype mice. Using a dose that resulted in maximal intestinal damage while avoiding acute deaths, we assessed early post-injury epithelial proliferation and determined that FR resulted in greater intestinal crypt cell proliferation at 3.5 days post-irradiation compared to SR (*34, 35*). To expand upon this previous work and investigate how this preserved epithelial proliferation impacts radiation-induced toxicity, we monitored the weight and survival of thirty-six male C57BL/6 mice after 14 Gy single-fraction whole-abdomen FR or SR (**Fig. 1A**). Post-treatment weight differences between FR and SR were temporally dependent (p < 0.001), with FR significantly improving weight recovery beginning at day 9 and extending until ten weeks post-RT (**Fig. 1B**). FR also resulted in improved overall survival that persisted for nearly 300 days after irradiation (p = 0.017) (**Fig. 1C**), which is near the expected lifespan of this mouse strain. Importantly, this greater survival with FR was linked to greater weight recovery at day 14, representing the end of the acute post-irradiation period (HR of SR vs. FR: 1.125, p = 0.002) (**Fig. 1D**). Thus, the intestinal epithelial sparing effects of FR manifest as greater overall survival involving a mechanism that likely begins with improved acute post-irradiation weight recovery.

**Figure 1.**
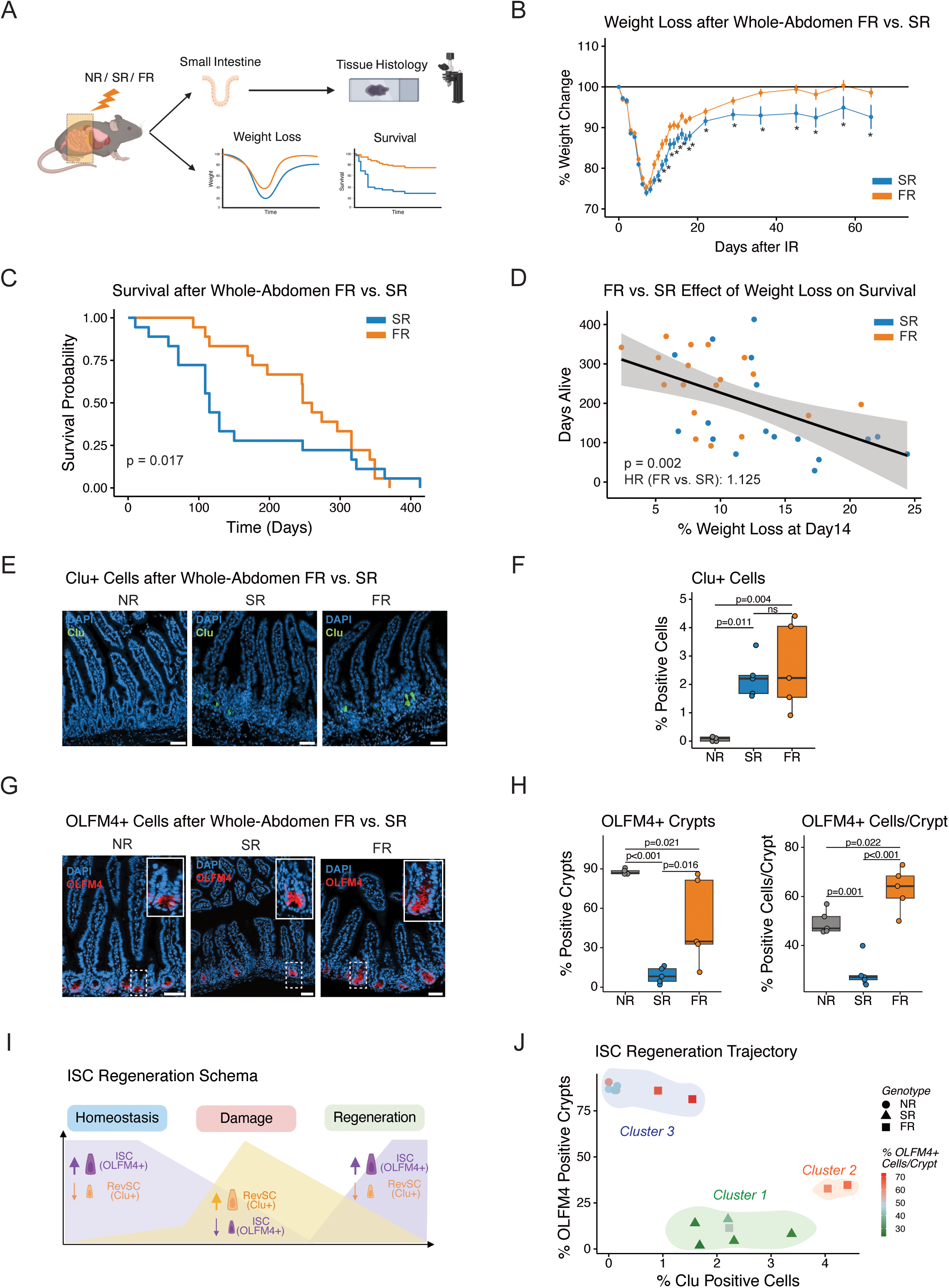
Enhanced crypt regeneration and survival benefit following whole-abdomen FR as compared to SR. **(A)** Schematic of experimental design after whole-abdomen (WA) irradiation. **(B)** Percentage of body weight change at specified time points following 14Gy WA irradiation with SR (n=18) or FR (n=18). The black line indicates the baseline. P-values at each time point were calculated using contrasts within a linear mixed-effects model. **(C)** Kaplan-Meier survival analysis of mice post-14 Gy WA SR (n=18) or FR (n=18). The Gehan-Breslow-Wilcoxon method was used to compare survival across treatment groups. **(D)** Correlation between percentage weight loss at day 14 and days alive after 14Gy WA SR or FR. Each point represents an individual mouse. The black line represents the linear regression line fitted to the data. Cox proportional-hazard models assessed the impact of weight loss and irradiation on survival. **(E)** Representative images of *Clu* RNA in situ hybridization on tissue sections from isolated small intestines non-irradiated (NR) or 3.5 days after SR or FR treatment. Scale bar = 50μm. **(F)** Percentage of *Clu*+ cells quantified from (E). The boxplot represents the median and interquartile range (IQR). Whiskers represent the highest and lowest values within 1.5 times the IQR. Each dot is the average of 15 tissue areas quantified per mouse. Statistical significance was calculated using one-way ANOVA followed by post-hoc Tukey’s HSD test. Insets highlight the number of OLFM4+ cells/crypt. Scale bars = 50µm. **(G)** Representative images of OLFM4 immunofluorescence on tissue sections from isolated NR small intestines or 3.5 days after SR or FR treatment. Scale bar = 50um. **(H)** Percentage OLFM4+ crypts and OLFM4+ cells per crypt quantified from (G). Boxplot represents median with IQR. Whiskers represent highest and lowest values within 1.5 times the IQR. Each dot is the average of 15 tissue areas quantified per mouse. Statistical significance calculated using one-way ANOVA followed by post-hoc Tukey’s HSD test. **(I)** Schematic of tissue regeneration phases following irradiation, including the dynamic expression of the intestinal stem cell (ISC) marker OLFM4 and revival stem cells (RevSC) marker *Clu*. **(J)** Regeneration trajectory based on percentage of OLFM4+ crypts and *Clu*+ cells in NR (circle), SR (triangle), and FR (square) groups. Color intensity represents the percentage of OLFM4+ cells per crypt.

To explore how FR might differentially impact epithelial cells and improve weight recovery and survival, we first investigated intrinsic differences in ISC survival after radiation-induced damage. *In vitro* irradiation of intestinal organoids, which is largely comprised of ISCs, with either 8 Gy single-fraction FR or SR resulted in equivalent depletion of ISCs, suggesting that the FLASH effect requires the intestinal microenvironment **(fig. S1A-C)**. Therefore, we characterized post-irradiation intestinal crypt regeneration *in vivo* by following ISC differentiation dynamics using immunofluorescence (IF) and RNA in situ hybridization at 3.5 days after SR or FR. Typically, radiation injury results in disruption of crypt architecture and rapid death of radiation-sensitive homeostatic ISCs that can be identified using either *Lgr5* or *Olfm4* stem cell markers. This radiation-induced death of *Olfm4*+ ISCs is compensated by emergence of *Clu*^+^ damage-induced revSCs that function to replenish *Olfm4*^+^ ISCs residing in intestinal crypts (*26*). Upon restoration of *Olfm4*^+^ ISCs to homeostatic levels, *Clu*^+^ revSCs promptly return to quiescence and become nearly undetectable. As expected, both SR and FR led to a decrease in OLFM4^+^ crypts that was accompanied by an increase in *Clu*^+^ cells per high-power field (**Fig. 1E-H**). Notably, when compared to SR, FR resulted in a higher number of OLFM4^+^ crypts (p = 0.016) and in more OLFM4^+^ cells per crypt that exceeded homeostatic levels (p < 0.001), consistent with ongoing regeneration. Plotting the percent of OLFM4^+^ crypts as a function of the percent of *Clu*^+^ cells for each treatment group summarized this regeneration trajectory (**Fig. 1I-J**). Here, cluster 1 captured the *Clu*^+^ revSCs formed after radiation-induced damage. Cluster 2 highlighted the conversion to and proliferation of OLFM4^+^ stem cells. Cluster 3 represented the *Clu*^+^ revSCs returning to quiescence and the reconstitution of the intestinal crypt, as well as homeostatic ISCs from non-irradiated controls. Overall, there was a greater proportion of FR versus SR samples in clusters 2 and 3 (p = 0.047). These data suggest that enhanced post-irradiation weight recovery with FR was associated with accelerated crypt regeneration coordinated by damage-induced revSCs.

### Temporal tissue and transcriptional changes associated with epithelial damage and regeneration after FLASH

To obtain a more detailed understanding of how FR could enhance intestinal regeneration compared to SR, we performed single-cell RNA-sequencing (scRNA-seq) on dissociated epithelial and lamina propria cells of small intestines at 2, 3.5, 10, and 20 days after 14 Gy whole-abdomen irradiation. Selected timepoints revolved around our previous crypt regeneration assay at day 3.5 post-irradiation showing significantly higher proliferating cells per crypt in intestines treated with FR versus SR (*34*). All timepoints included epithelial and lamina propria cells except for day 3.5, which included only the epithelial fraction (**Fig. 2A**). Integration of 33 samples resulted in 132,246 high-quality cells that were embedded into low dimensional UMAP space. Clustering followed by annotation using cell type specific gene markers revealed established small intestinal cell populations that spanned immune (62%), epithelial (29%), stromal (9%), and neural (<1%) cell types. Further sub-clustering resulted in 46 cell types with distinct transcriptional profiles (**Fig. 2B-C**, **fig. S2A-B, and Data File S1)**. One cluster labeled as “Regenerating Crypt Cells” (regCCs) was unique to irradiated samples (**Fig. 2D**). These regCCs were marked by high expression of a previously described regenerative crypt gene signature and were separate from the *Lgr5*+ and *Olfm4*+ intestinal stem cells **(fig. S2-D)** (*26*). Substructure analysis within this cluster revealed quiescent revSCs and a proliferative regenerative crypt base columnar (CBC) cell cluster that may represent revSCs transitioning from quiescence (SSC2a) **(fig. S2E)** (*26, 36*). After irradiation and death of the majority of differentiated epithelial cells, these regCCs constituted most of the intestinal crypt but then declined over time as other CBC cells regenerated (**Fig. 2D-E**). Thus, these data represent a comprehensive representation of intestinal biology that captures the transcriptomic temporal dynamics of tissue regeneration after FR and SR.

**Figure 2.**
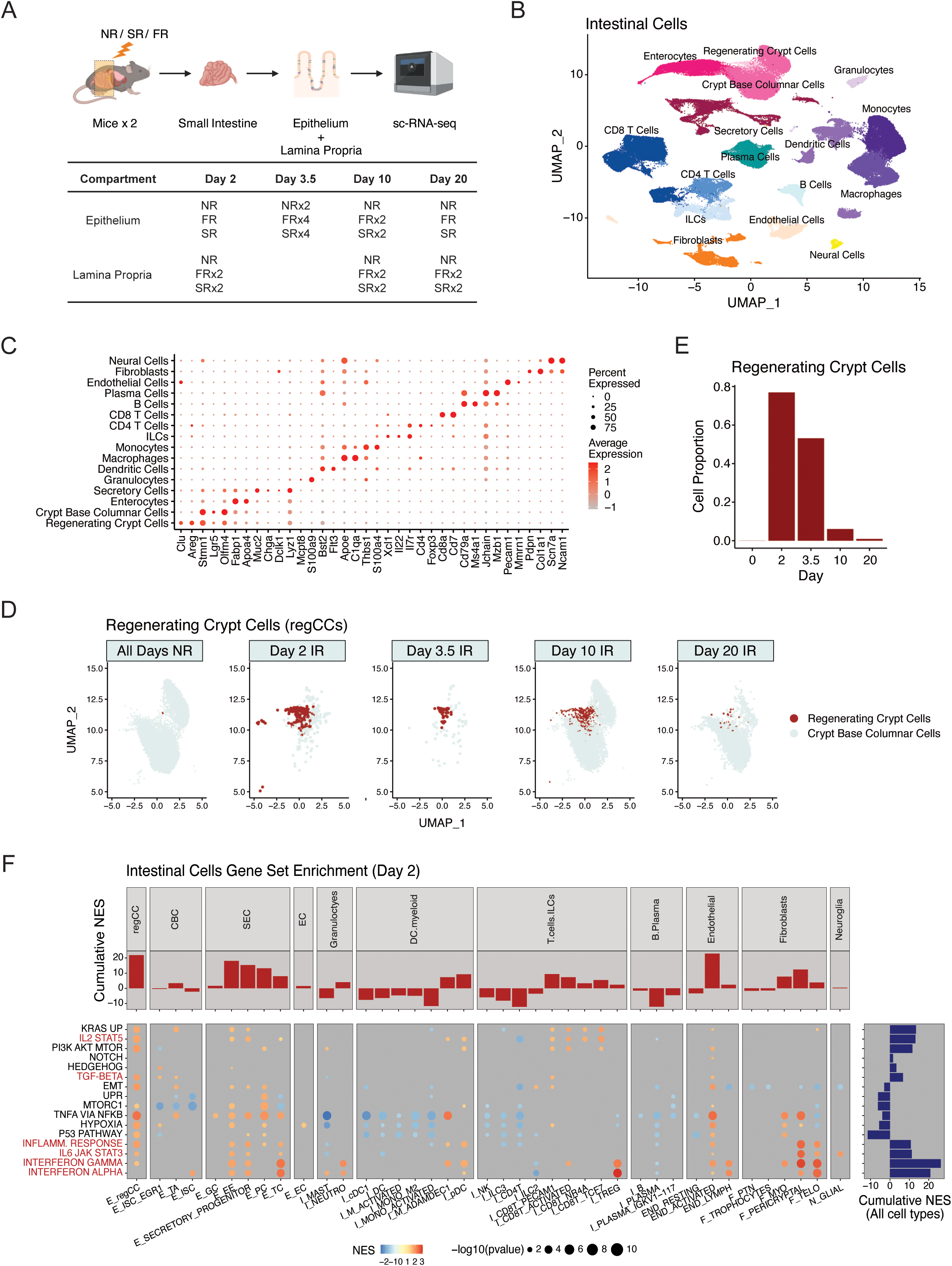
Time course scRNA-sequencing captures intestinal regeneration and uncovers inflammatory signaling pathways associated with FLASH RT. **(A)** Schematic illustrating the process of harvesting small intestines and isolating epithelium and lamina propria fractions for single-cell RNA sequencing (scRNA-seq) at specified time points. **(B)** Uniform Manifold Approximation and Projection (UMAP) plot showing all epithelial and lamina propria cell clusters captured at all time points, color-coded by general cell type. **(C)** Dot plot displaying selected lineage-specific gene markers across all cell clusters. The color intensity represents the average expression, while dot size indicates the percentage of cells expressing each gene. **(D)** UMAP plots of regenerating crypt cells (regCCs, red) and crypt base columnar cells (CBCs, light gray) at selected time points following RT. **(E)** Bar plot of the proportion of regCCs at selected time points following RT. **(F)** Gene set enrichment analyses (GSEA) on gene sets from the Hallmark Collection across all cell clusters, comparing FR to SR samples at day 2 post-RT. The color intensity represents the Normalized Enrichment Score (NES), and dot size represents p-value. Growth and inflammatory signaling pathways upregulated in FR compared to SR are highlighted.

### FLASH is associated with increased inflammatory signaling in regenerative stem cells and other distinct cell types

We next sought to understand the transcriptomic changes resulting from FR compared to SR by first performing pathway-level analysis using gene set enrichment analysis (GSEA) on the Hallmark Gene Set Collection. This revealed that FR led to significant enrichment of cell cycle and proliferation-related gene sets within regCCs at day 3.5 post-irradiation, consistent with the enhanced EdU incorporation observed with FR *in vivo* **(fig. S2F)** (*34*). To better understand the biology of the proliferative transition between days 2 and 3.5, we performed GSEA on signaling-related gene sets across all cell types, comparing FR to SR at day 2 post-treatment. Overall changes in signaling for each cell type and for each signaling pathway were captured by summing the normalized enrichment scores comparing FR versus SR either column-wise or row-wise, respectively (**Fig. 2F**). This revealed that regCCs, followed by endothelial cells, demonstrated the greatest overall increase in signaling activity enrichment (**Fig. 2F**, **red bar plots, top)**. Individual signaling pathways more enriched in regCCs after FR versus SR included TGF-β signaling, which has been reported to induce revSCs two days post-irradiation, in addition to KRAS and mTOR signaling, which is consistent with the greater proliferation after FR at 3.5 days post-irradiation (*28*). Examination of signaling pathways with large overall differences across cell types after FR compared to SR highlighted type I (IFN-I) and type II (IFNG) IFN signaling, and to a lesser extent, JAK-STAT activity and JAK-regulated cytokine pathways (e.g., IL-2 and IL-6) (**Fig. 2F**, **navy bar plots, right margin)**. Among cell types with the greatest enrichment for IFN and JAK-STAT pathways after FR included mesenchymal populations such as pericryptal fibroblasts and telocytes that both support ISCs through cytokine and growth factor signaling (*37, 38*).

Collectively, these findings outline tissue-level and cell type-specific signaling changes that differ between FR and SR and may relate to how revSCs undergo accelerated epithelial regeneration. Notable pathways include TGF-β, which can induce revSCs, and mitogenic pathways that promote revSC proliferation. Other pathways contributing to differences in FR versus SR include IFN signaling, which demonstrated the greatest increase in mesenchymal subsets known to support ISCs.

### Increased TGF-β signaling in revSCs after FLASH is associated with greater myeloid cell infiltration

We surmised that the tissue and cell type-specific signaling changes after FR and SR coordinate key cell-cell communication events involving regCCs, including revSCs, that may contribute to the FLASH effect. To begin investigating this notion, we used cell-cell interaction analyses to examine afferent signaling pathways to regCCs two days after FR versus SR **(fig. S3A)**. Consistent with GSEA and EdU incorporation studies, this revealed that the top afferent pathways to regCCs augmented by FR included TGF-β signaling, an inducer of revSCs, and FGF signaling, a mitogenic pathway. Examining potential cellular sources for afferent TGF-β signals into regCCs that were greater with FR versus SR nominated monocyte and macrophage populations (**Fig. 3A**), which is consistent with previous reports (*28*). Indeed, after irradiation the *Tgfbr2* receptor and the *Tgfb1* cytokine ligand were identified by receptor-ligand interaction analysis (**Fig. 3B**), and both genes were highly and specifically expressed in regCCs and in macrophages and infiltrating monocytes, respectively **(fig. S3B-C)**. Thus, elevated myeloid-derived TGF-β signaling, a known source and inductive signal for revSCs, may contribute to enhanced immune-mediated crypt regeneration kinetics after FR.

**Figure 3.**
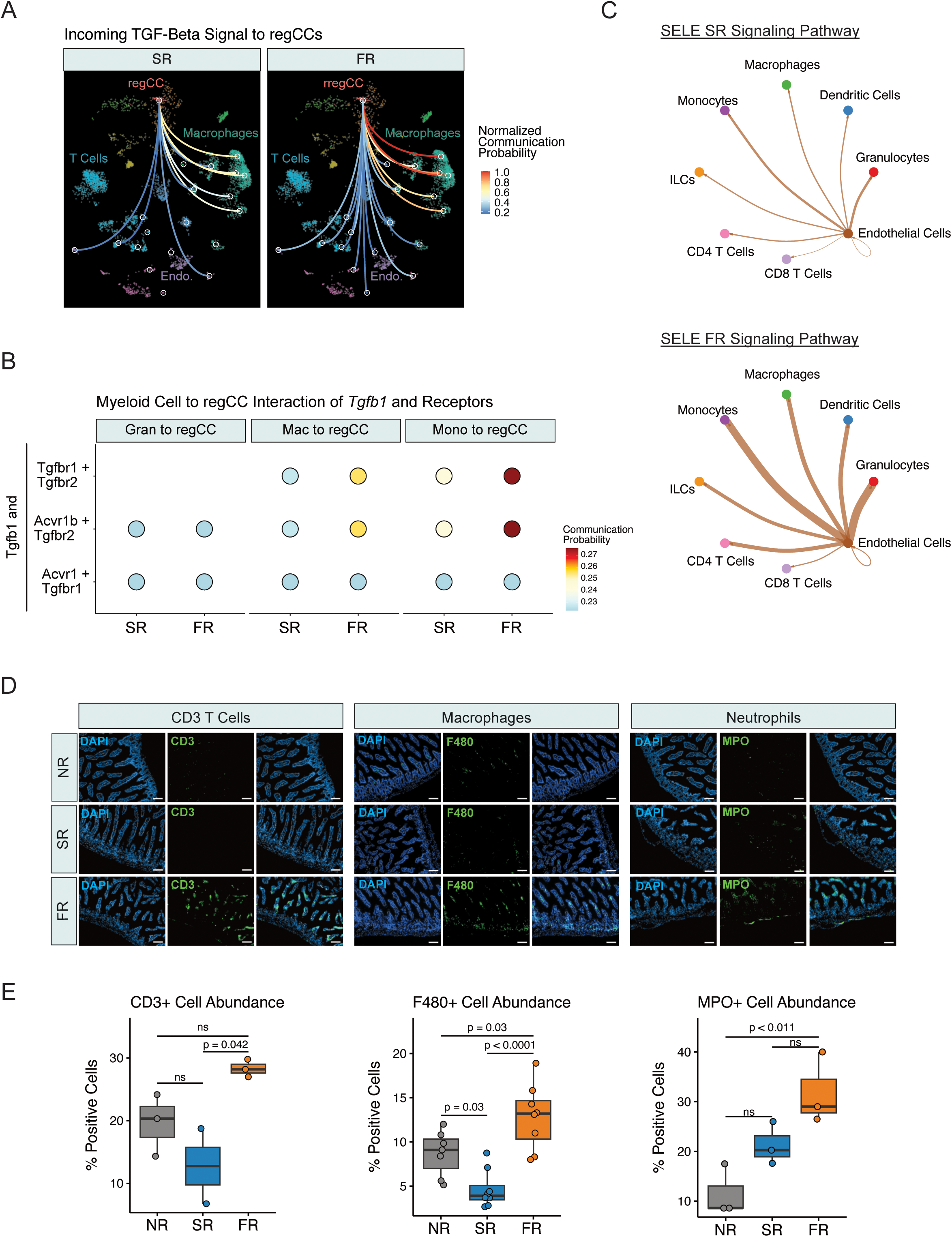
Increased crypt regeneration in FLASH treatment is associated with higher TGF-β-induced immune cell infiltration. **(A)** Network analysis showing cell communication enrichment for TGF-β signaling from stromal cells to regenerating crypt cells (regCC) 2 days post-SR or FR. **(B)** Bubble chart illustrating TGF-β ligand-receptor pair signaling from myeloid subtypes (granulocytes (Gran), macrophages (Mac), and monocytes (Mono)) to regCCs, calculated by CellChat. Dot color reflects communication probabilities, and dot size represents computed p-values. **(C)** Interaction analysis for E-selectin signaling pathway between endothelial cells and other cell types for SR (top) and FR (bottom). Line thickness indicates the strength of interaction for each cell type. **(D)** Representative immunofluorescent images from isolated small intestines stained for CD3 (T cells), F4/80 (macrophages), and MPO (neutrophils) from non-irradiated (NR) or two days after SR or FR. Magnification, ×10. Scale bars, 100 μm. **(E)** Percentage of CD3, F4/80, or MPO-positive cells per condition from (D). Boxplot represents median with interquartile range (IQR). Whiskers represent highest and lowest values within 1.5 times the IQR. Each dot is the average of 10 tissue areas quantified per mouse (n=3 for CD3 and MPO, n=7 for F4/80, biologically independent samples per group). Statistical significance calculated using one-way ANOVA followed by post-hoc Tukey’s HSD test.

Next, we sought to investigate how myeloid-derived TGF-β signaling in revSCs might be augmented after FR compared to SR. Besides regCCs, endothelial cells exhibited among the largest enrichment of multiple signaling pathways (**Fig. 2F**). Moreover, cell-cell interaction analysis identified E-selectin as one of the most upregulated signaling pathways after FR compared to SR. Investigation of sender-receiver pairs predicted to communicate through E-selectin signaling revealed significantly more interactions between activated endothelial cells and immune cells after FR versus SR, particularly through SELE-CD44 signaling (**Fig. 3C**, **fig. S3D)**. Based on these findings, we used IF staining of immune cell populations two days after RT to explore if immune cell infiltration might be altered by FR compared to SR. This revealed that macrophage infiltration into the regenerating intestines was indeed significantly greater after FR compared to SR (**Fig. 3D-E**). An increase in T cell but not neutrophil infiltration was also observed following FR.

Together, these findings suggest that FR results in greater myeloid cell infiltration, possibly through an effect on endothelial cell signaling. This greater infiltration is associated with enhanced TGF-β signaling in regCCs after FR, consistent with myeloid cells being a primary source of TGF-β important for revSC induction and intestinal regeneration after radiation-induced damage (*28*). Thus, enhanced myeloid infiltration and inductive signals by TGF-β may contribute to accelerated intestinal regeneration after FR.

### FLASH drives enhanced interferon signaling in mesenchymal cells that support intestinal stem cells

IFN signaling was one of the most enriched pathways after FR compared to SR (**Fig. 2F**). Closer examination of IFN-I and IFNG pathway enrichment across the epithelium and lamina propria highlighted that although IFN signaling was significantly enriched in both compartments, the lamina propria showed the strongest contribution **(fig. S4A)**. Within the lamina propria, pericryptal fibroblasts and telocytes were confirmed to have the greatest enrichment (**Fig. 2F**, **Fig. 4A-B**). This was additionally corroborated by qRT-PCR for several IFN-stimulated genes (ISGs) using dissociated intestinal lamina propria tissue from mice after SR or FR. At day 1 post-irradiation, there was equal ISG induction between the FR and SR groups. However, by days 2 and 3 post-irradiation, several ISGs showed a trend towards increased expression in the FR compared to the SR group with aggregation of these fold changes consistent with significantly increased IFN pathway activation (**Fig. 4C**). Thus, these data suggest that compared to SR, FR drives greater IFN signaling in pericryptal fibroblasts and telocytes *in vivo*.

**Figure 4.**
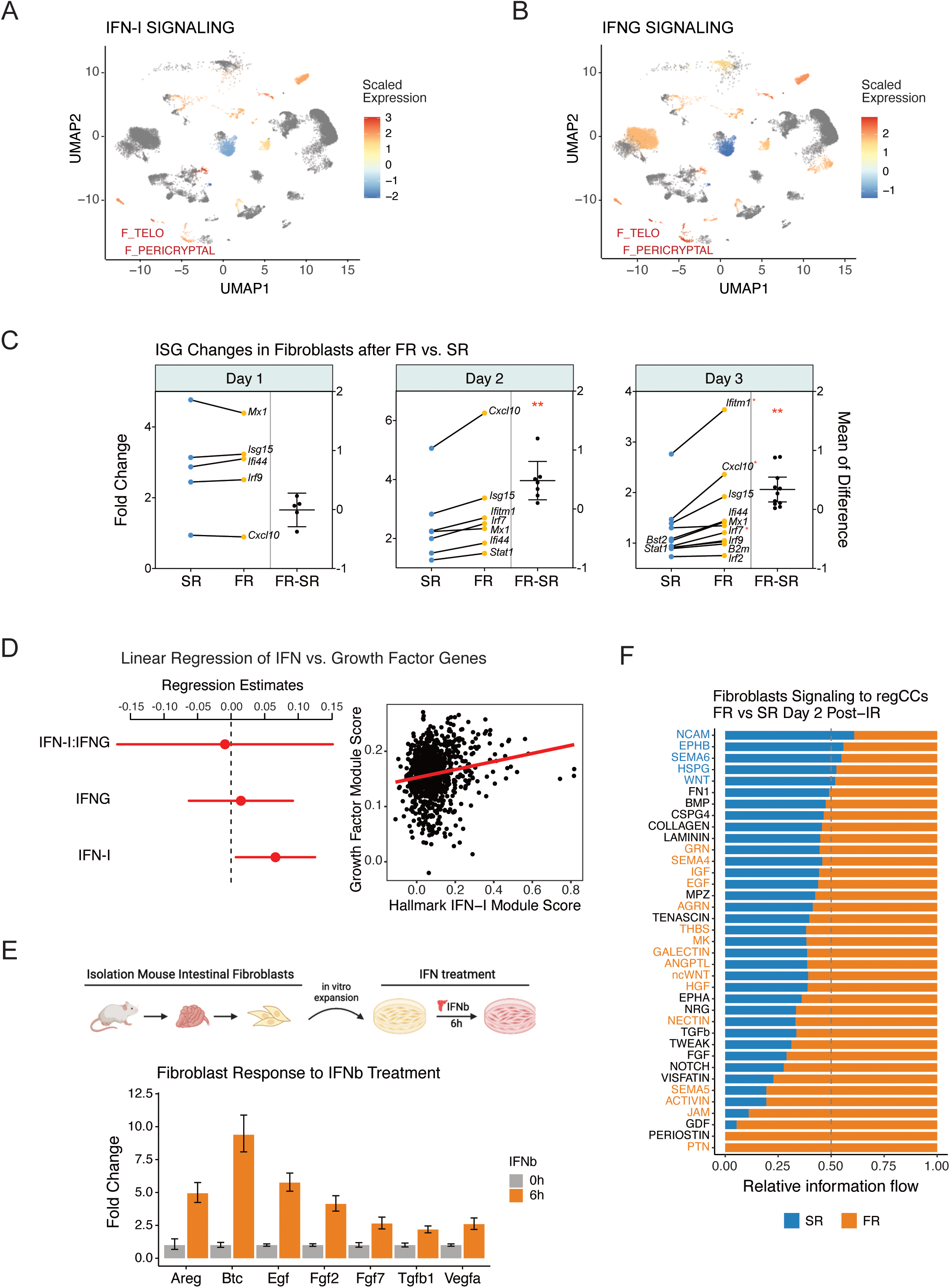
FR induces higher Type I IFN signaling in lamina propria cells and is associated with enhanced crypt regeneration. **(A, B)** UMAP plots displaying the average expression of Type I and II IFN gene signature expression across all cell types and time points. **(C)** Fold change expression of individual interferon-stimulated genes in lamina propria cells isolated from small intestines at indicated time points following SR or FR as normalized to NR. **(D)** Regression analyses between Reactome IFN signaling pathways and growth factor module score in pericryptal fibroblasts and telocytes 2 days post-RT. **(E)** Schematic of intestinal fibroblast isolation, in vitro expansion, and treatment with IFN-β. Fold change in expression of indicated genes on intestinal primary fibroblasts at 0 and 6 hours following IFN-β treatment. **(F)** Stacked bar plot comparing the inferred molecular flow of information from fibroblasts to regCCs between FR and SR two days after RT. Pathways labeled in blue indicate significantly greater information flow in SR, while orange labels indicate greater flow in FR.

IFN-I signaling has been previously linked to tissue regeneration by EGF and FGF induction, amongst other growth factors, in lamina propria cells (*39–41*). However, distinguishing between the effects of IFN-I versus IFNG is challenging given the correlated expression of the ISGs that have been ascribed to each IFN pathway. To examine whether IFN-I independently correlates with growth factor activity in fibroblasts while controlling for its correlation for genes regulated by IFNG signaling, we utilized linear regression that considered both IFN-I and IFNG gene set expression scores, or activity. Indeed, this revealed that IFN-I, but not IFNG activity, was independently and positively correlated with growth factor expression in pericryptal fibroblasts and telocytes after irradiation *in vivo* (**Fig. 4D**, **fig. S4B-C)**. Consistent with this, *in vitro* treatment of primary intestinal fibroblasts for six hours with IFN-I (IFN-β) corroborated the ability of IFN-I to transcriptionally induce FGF and EGF family genes, as well as *Vegfa*, *Tgfb1*, and other related cytokines (**Fig. 4E**). Cell-cell communication analysis of irradiated intestinal cells two days after FR versus SR predicted a significant likelihood that fibroblasts and telocytes are likely the main sources of FGF and EGF growth factor signaling between fibroblasts and regCCs (**Fig. 4F**, **fig. S4D-E)**. Collectively, these results indicate that greater acute elevations in IFN-I signaling in FR compared to SR may promote fibroblast production of FGF and EGF growth factors that support regCC proliferation and crypt regeneration.

### Type I IFN signaling is specifically required for the accelerated epithelial regeneration and decreased morbidity resulting from FLASH

In disease states such as pathogen infection and cancer, IFN-I signaling can have complex and even opposing roles in intestinal differentiation and regeneration after injury (*42, 43*). Thus, we sought to extend our analysis to examine what pathways in addition to growth factor activity in fibroblasts are independently associated with IFN-I signaling. Moreover, we specifically focused on pathways whereby this association might differ between FR versus SR after controlling for effects of IFNG pathway activity (see Methods). This analysis revealed multiple pathways in distinct cell types that correlated with IFN-I signaling but in a manner that differed between FR and SR (**Fig. 5A**). In regCCs for example, differences in how IFN-I activity was correlated with p53-related genes was driven by a greater proportion of regCCs with increased p53 pathway expression after FR versus SR (87.3% vs. 67.6%), which is consistent with our findings from GSEA (**Fig. 5A**, **Fig. 2F**). The greater proportion of cells with high p53 activity and its altered correlation with IFN-signaling with FR compared to SR were notable because we have recently shown that p53 promotes intestinal regeneration after radiation through induction of revSCs (*27*). In pericryptal fibroblasts and telocytes, expression of multiple FGF family members was predicted to positively associate with IFN-I activity after FR but negatively correlated after SR, findings consistent with the greater intestinal crypt cell proliferation after FR compared to SR (**Fig. 5B**). Together, these results suggest that the impact of IFN-I signaling on pathways that include those that influence intestinal regeneration may differ after FR compared to SR.

**Figure 5.**
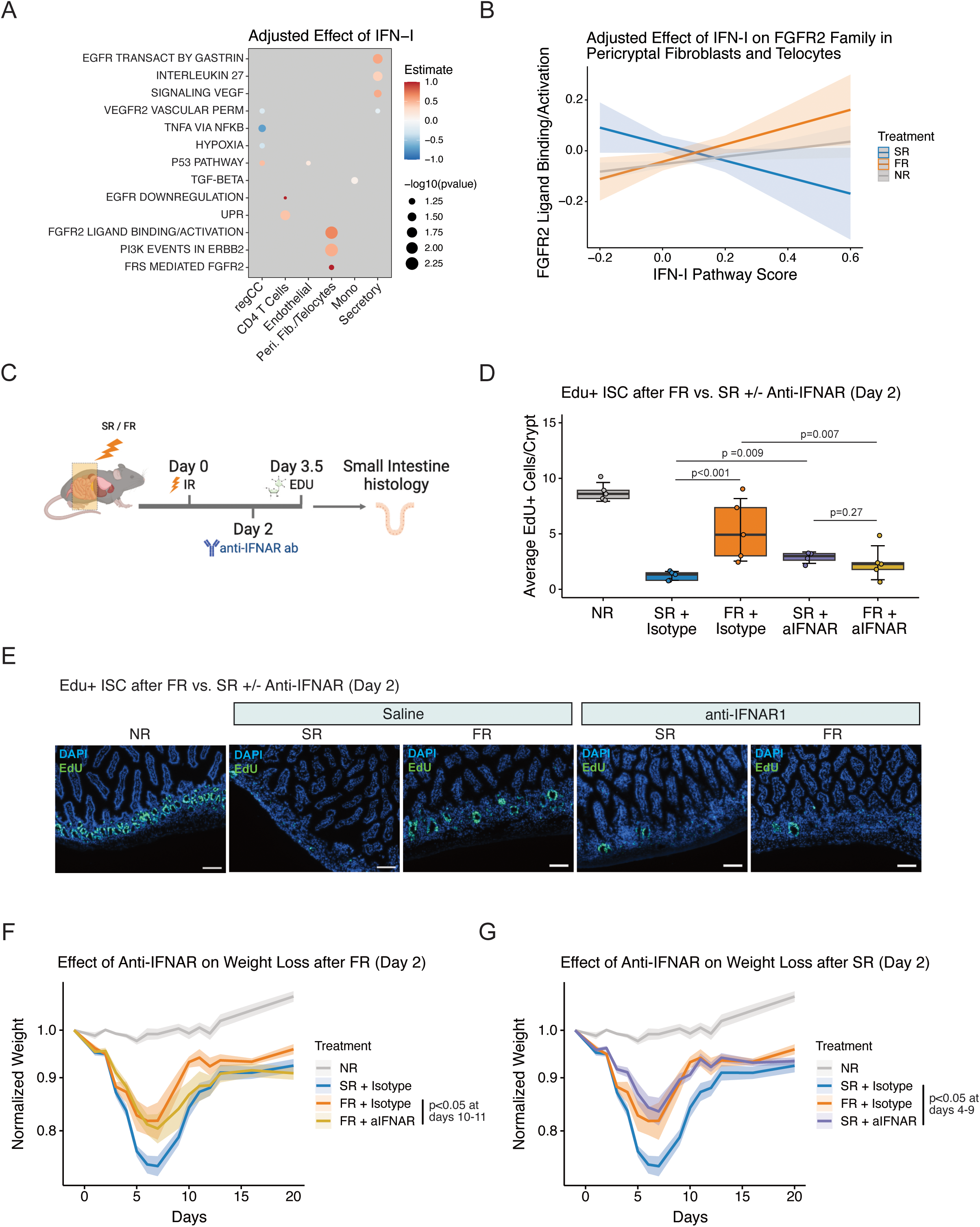
FR-associated regeneration effect is abolished following type I IFN blockade. **(A)** Adjusted effect size of type I IFN signaling on various pathways involved in tissue regeneration. Adjusted effect size was determined from regression analyses, controlling for contributions from IFNG signaling. Dot color signifies effect size (regression coefficient), while dot size represents p-value. **(B)** Effect of type I IFN signaling on FGFR2 signaling, stratified across treatment modality. **(C)** Schematic representation of whole-abdomen RT and follow-up assays. Anti-IFNAR antibody was administered on Day 2 post-RT. **(D)** Average number of EdU+ cells per crypt in the intestines of mice 3.5 days after no irradiation (NR), standard radiation (SR), or FLASH radiation (FR) with or without IFNAR1 blocking antibody treatment. Boxplot represents interquartile range (IQR) with median indicated. Whiskers represent highest and lowest values within 1.5 times the IQR. Each dot is the average of 10 tissue areas quantified per mouse (n=5 biologically independent samples per group). Statistical significance was calculated using two-factor ANOVA testing followed by estimated marginal means. **(E)** Representative immunofluorescent images from isolated small intestines stained for EdU at Day 3.5 after NR, SR, or FR with or without IFNAR1 blocking antibody treatment. Magnification, ×10; Scale bars, 100 μm. **(F-G)** Body weight loss normalized to Day 0 for each indicated time point and treatment condition. Statistical significance was calculated using linear mixed-effect models with post-hoc contrasts.

To examine if IFN-I signaling regulates tissue regeneration and morbidity after radiation-induced damage and whether it can specifically contribute to the FLASH effect, we treated mice with an anti-IFNAR1 blocking antibody at two days or 12 hours after FR or SR (**Fig. 5C**, **fig. S5A)**. At both day 2 and 12 hours post-irradiation, anti-IFNAR1 decreased EdU incorporation in intestinal crypt cells compared to isotype control, essentially abrogating the improved crypt cell proliferation observed with FR (p = 0.007 and p = 0.003, respectively) (**Fig. 5D-E**, **fig. S5B-C)**. In distinct opposition, anti-IFNAR1 increased the degree of EdU incorporation in crypt cells after SR, suggesting that rather than promoting intestinal regeneration as observed after FR, IFN-I signaling after SR antagonized early regeneration events. Consistent with this, examination of weight recovery after FR revealed that IFNAR1 blockade also abrogated the improved weight recovery afforded by FR compared to SR, but primarily at days 10-11 post-irradiation (p < 0.05). No impact of anti-IFNAR1 on weight recovery was observed at days 4-9 (**Fig. 5F**), suggesting that anti-IFNAR1 specifically interfered with the later phases of weight recovery accelerated by FR. In contrast to these effects after FR, anti-IFNAR1 treatment after SR improved the early weight loss at days 4-11 (p < 0.05) that is typically worse with SR (**Fig. 5G**), resulting in overall weight recovery similar to FR.

In total, these divergent effects of anti-IFNAR1 on tissue regeneration and weight recovery after FR as opposed to SR mirror the variable associations between IFN-I signaling and pathways participating in intestinal regeneration. Namely, IFN-I signaling is required for enhanced intestinal regeneration and weight recovery specifically after FR, while after SR, IFN-I signaling rapidly drives detrimental effects that worsens morbidity. These opposing effects are associated with differences between FR and SR in how IFN-I signaling correlates with pathways controlling p53 induction in regCCs and with FGF expression in ISC-supporting fibroblasts.

## DISCUSSION

Unraveling the biological foundations of the FLASH effect could facilitate its clinical adoption and offer new ways to decrease treatment toxicities potentially across various cancer treatments. Here, we found that the tissue sparing effect of FR in the intestines of mice after whole-abdomen radiation was linked to accelerated differentiation of ISCs, including the revSC damage-induced stem cell population, and to the elaboration of a tissue regenerative inflammatory milieu characterized by TGF-β and IFN-I signaling. TGF-β signaling, a known inducer of revSCs, appeared to increase more with FR due to greater immune cell infiltration of myeloid cells producing TGF-β and due to higher levels of its receptor *Tgfbr2* expressed by revSCs. IFN-I signaling, a less understood pathway implicated in intestinal epithelial regeneration, increased in the lamina propria compartment to a greater degree after FR, resulting in evidence for enhanced growth factor induction in fibroblast subsets known to support intestinal epithelial proliferation. Taken together, these findings suggest that a cellular network of immune cells, damage-induced stem cells, and stromal cells coordinated by IFN-I signaling contribute to the FLASH effect. Unexpectedly, not only is IFN-I signaling noncontributory to tissue regeneration after SR, in this context it appears to serve a dichotomous role by promoting the increased toxicity observed after SR compared to FR.

The balance between intestinal stem cell differentiation and mechanisms that preserve the stem cell pool is critical for tissue homeostasis and for ensuring rapid tissue regeneration after injury; moreover, its imbalance may have roles in disease pathology. As such, differences in how FR versus SR stimulate regulatory programs that control the balance between stem cell differentiation vs. self-renewal could also underlie the FLASH effect. Recent work has described competing effects between TGF-β-driven YAP signaling and EGFR-activated MAPK/PI3K activity that can induce a *Clu*+ revSC quiescent phenotype or incite a proliferative response, respectively (*44*). Indeed, it can be speculated that the increased proliferative shift in regCCs from day 2 to day 3.5 after FR compared to SR may represent differences in the balance between such regulatory programs – an FR-mediated increase in myeloid-derived TGF-β that drives revSC induction and an increase in subsequent fibroblast-derived FGF that promotes OLFM4+ crypt cell proliferation and expansion. Thus, the greater proportion of intestinal stem/progenitor cells that we find at later states in the regenerative trajectory after FR versus SR attests to accelerated crypt renewal, likely spurred by early increases in both TGF-β and FGF signaling that characterizes FR.

A central role for IFN-I signaling in coordinating a cellular network that contributes to the FLASH effect is a key yet also perplexing finding of our study. Our data suggests that FR results in both quantitative and qualitative differences in IFN-I signaling compared to SR that are associated with the improved morbidity and mortality. Besides apparent quantitative changes such as the greater magnitude of IFN-I pathway gene induction after FR versus SR in mesenchymal cells, our findings also suggest qualitative differences in IFN-I signaling with FR compared to SR. Linear modeling of transcriptomic data suggests that after controlling for IFNG signaling, increases in IFN-I signaling activity positively correlates with expression of FGF family genes in pericryptal fibroblasts and teleocytes after FR but negatively correlates after SR. This suggests that in contrast to FR, IFN-I signaling after SR may limit the elaboration of FGF and perhaps other growth factors that support regCC proliferation and tissue regeneration. Indeed, our experiments with anti-IFNAR1 are consistent with the interpretation of this statistical modeling – namely, IFN-I signaling after SR interferes with regeneration to worsen toxicity while IFN-I signaling after FR promotes tissue repair to decrease morbidity.

Although additional investigation is needed to understand how FR results in altered IFN-I signaling that affects tissue repair conversely to SR, such findings that IFN can have context-dependent and dichotomous roles in intestinal injury and repair have been observed previously. In mouse models of radiation-or DSS-induced intestinal injury, IFN-I has been reported to promote intestinal repair by enhancing stem cell proliferation through cGAS/STING, a DNA pattern recognition receptor (PRR) pathway (*45*). In models of immune-mediated damage of intestinal tissue from GVHD, activation of IFN through RNA sensing PRRs can also promote repair (*46*). However, although these favorable properties of IFN-I can acutely enhance tissue regeneration by driving progenitor cell proliferation and differentiation, continuous activation can also interfere with tissue homeostasis. For example, persistent IFN-I signaling from chronic viral infection or prolonged activation of RNA-sensing PRRs can unremittingly drive secretory-cell differentiation and exhaust intestinal stem cell pools (*30*). Likewise, other IFNs that can be co-regulated with IFN-I can also negatively impact intestinal homeostasis and injury repair. IFNG can promote intestinal injury directly by causing epithelial cell death or indirectly through T cells (*47*). Both IFNG and IFN-I can synergize with TNF-α to foster intestinal epithelial cell death and retard regeneration (*48*). Interestingly, our findings show that TNF-α is another pathway that exhibits an attenuated correlation with IFN-I signaling after FR compared to SR (Fig. 5A), raising the possibility that FR might yield less cooperativity between IFN-I and TNF-α at promoting epithelial cell death, contributing to the FLASH effect.

Together, our results suggest that the pleiotropic effects of IFN signaling on intestinal regeneration and homeostasis may contribute to the normal tissue sparing effect of FR. Future studies are needed to determine if differences in how this pleiotropy is manifested after FR compared to SR can mechanistically explain the FLASH effect, whether these IFN-related mechanisms are similar across other tissues where the FLASH effect has been observed (*5–9*), and how other proposed mechanisms for the FLASH effect such as hypoxia might relate. This deeper understanding may also have implications for improving toxicities from other cancer therapy modalities as well.

## MATERIAL AND METHODS

### Study Design

For the in vivo survival and weight loss studies, 18 mice were allocated to each RT treatment modality (SR or FR). Additionally, 5 non-irradiated mice served as controls (NR group). Throughout the study period, animals were monitored until they reached the experimental endpoint or exhibited weight loss >25% of their initial body weight. No animals or data points were excluded from the analysis. This study lasted over a year, and it was done once. For the body weight loss and EdU incorporation studies involving anti-IFNAR antibody treatment, a total of 5 mice were used for each condition (NR, SR, and FR). These studies have been conducted a total of 4 times (twice at 2 days post-treatment and twice for 12h post-treatment). One representative example of each is shown in this manuscript. For the EdU incorporation experiments, the immunofluorescence staining of immune cells and OLFM4+ cells, and the RNA in situ hybridization of *Clu* mRNA, a total of five mice per condition were evaluated. Immunofluorescence and RNA in situ hybridization experiments were done once. Animal weight loss measurement and microscope image acquisition were performed by a blinded researcher. For the single-cell RNA sequencing experiments, single cells were pooled from 2 animals for each condition and time point.

### In Vivo Mouse Studies

Nine-to eleven-week-old male C57BL/6J mice were purchased from Jackson Laboratory (Bar Harbor, ME, USA). Animals were maintained in the University of Pennsylvania Association for Assessment and Accreditation of Laboratory Animals Care (AAALAC)-accredited animal facilities. All experimental procedures were conducted in accordance with protocols approved by the Institutional Animal Care and Use Committee. Mice were checked daily and euthanized upon onset of severe morbidity or apparent weight loss >20%. Mice were housed in 12:12 light–dark cycles with temperatures of ∼18–23 °C and 40–60% humidity. For the proton irradiation studies, 2% isoflurane in medical air was used to anesthetize mice. Mice were euthanized by CO_2_ asphyxiation.

### Proton Irradiation Experiments

For *in vitro* irradiation, intestinal organoids were plated and grown for two to three days in 12-well plates prior to irradiation. Culture media was removed before 8 Gy FR or SR and freshly added back immediately after irradiation. For *in vivo* irradiation, male mice were randomly assigned to either 14 Gy [SR: 0.82 ± 0.1, FR: 125.3 ± 4.8] whole-abdomen FR or SR as previously described, and intestinal segments were harvested at 2, 3.5, 10 or 20 days post-irradiation (*34*).

Irradiation was performed on a fixed angle research beam line using a 230 MeV (range ∼32 g/cm^2^) proton beam generated by an IBA Proteus Plus Cyclotron (Louvain-La-Neuve, Belgium) collimated to a field size of 2 x 2 cm^2^ for whole-abdomen irradiation or 26 mm diameter for cell irradiation. Field uniformity and alignment were verified prior to each experiment using EBT3 Gafchromic film (Ashland, Bridgewater, NJ, USA). Dose measurements were performed with a calibrated NIST-Traceable Advanced Markus Chamber (PTW, Freiburg, Germany), following the International Atomic Energy Agency Code of Practice TRS-398 (*49*). Online dosimetry was performed with a cross-calibrated thin window Bragg peak chamber (Type 34070, PTW, Freiburg, Germany). Beam delivery structure was collected with a 1 GHz oscilloscope, enabling dose rate determination.

### Small Intestine Cell Isolation

Intestinal epithelial cell isolation was performed as described previously (*50*). In brief, small intestines were removed, and around 15 cm of the jejunum were isolated, washed with ice-cold 1X HBSS (Cytiva, #SH3058801), opened longitudinally, and divided into three pieces of roughly equal length. To obtain the epithelial cells, intestinal fragments were incubated in epithelial cell solution (10 mM EDTA [Invitrogen, #15575020]-HBSS, 10 mM HEPES [Invitrogen, #15630080], 100 U/ml Pen-Strep [Invitrogen, #15140122], and 2% FBS [Life Technologies, #26140079]) for 15 min at 37°C in continuous gentle shaking and then for 10 min on ice. Tissue fragments were then transferred into fresh ice-cold HBSS and vigorously shaken to isolate the epithelial fraction. Intestinal crypts were recovered by filtering the tissue suspension through a 70-μm-mesh-cell-strainer. Three consecutive epithelial fractions were collected, pooled, and centrifugated for 10 minutes at 500 g. Epithelial cell pellets were resuspended in pre-warmed TrypLE Select Enzyme (Thermo Fisher, #12563011) and incubated for 5 min at 37°C. Single cells were obtained by continuous pipetting for 15-20 minutes at room temperature. TrypLE was quenched by the addition of 100% FBS, and cells were washed twice with HBSS and finally filtered through a 40-μm-mesh-cell-strainer. To isolate the stromal cells from the lamina propria (LP), the remaining intestinal segments were incubated in LP digestion mix (RPMI1640, 100 U/ml Pen-Strep, 10 mM HEPES, and 2% FBS freshly supplemented with 100 μg/mL of Liberase TM [Roche, #5401127001] and 100 μg/mL of DNase I [Zymo Research, #E1010]), at 37°C for 30 minutes. Single cells were obtained by continuous pipetting until complete dissociation of the tissue fragments. The LP enzymatic dissociation was quenched by the addition of 100% FBS and 80 μL of 0.5 M EDTA. Finally, the cell suspension was washed twice with HBSS and filtered through a 40-μm-mesh-cell-strainer. Both epithelial and LP single-cell suspensions were treated with ACK Lysis buffer (Quality Bio, #118156721) for 5 minutes at room temperature to lyse red blood cells.

### Flow Cytometry Cell Sorting and 10X Single-Cell RNA Seq Processing

Epithelial and LP cell suspensions were incubated with 1μg/mL FxCycle Violet Stain (Thermo Scientific, #F10347) for 5 minutes prior to flow cytometry. For each sample, between 200K and 400K single and alive cells were sorted in Advanced DMEM/F-12 (Thermo Fisher, #12-634-028) supplemented with 10% FBS, washed twice with PBS, and counted using an automated cell counter. 10K cells per sample were used for single-cell 3’-RNA sequencing following the manufacturer protocol using the Chromium Next GEM single-cell 3’ v3.1 (dual Index) kit (10x Genomics, #PN-100268). A total of 33 samples were processed, and cDNA quality control was assessed by capillary electrophoresis (Bioanalyzer, Agilent) before library preparation and sequencing on NovaSeq 2500 (Illumina).

### Single-Cell RNA-Sequencing Data Processing and Integration

The scRNA-seq data was processed using the CellRanger pipeline (10x Genomics) to demultiplex the FASTQ reads, align them to the mm10 Mouse genome, and generate gene-barcode expression matrices. Quality control of these expression matrices were conducted using the *Seurat* R package to retain cells with greater than 500 unique UMIs and a proportion of counts in mitochondrial genes less than 15%. Additionally, only genes with non-zero counts in more than ten cells were kept, and doublet removal was performed using *scDblFinder*. Dataset integration was conducted using *scVI* followed by *scANVI* both with default parameters. Briefly, unsupervised clustering, cluster annotation with marker gene expression, and subclustering were conducted after initial *scVI* integration. Two clusters with significant B-cell contamination and one cluster with red blood cell features were removed. Semi-supervised integration with *scANVI* was then performed to enhance bio-conservation without sacrificing significant batch correction power. Identification of specific gene markers for all clusters was calculated using *Seurat’s FindAllMarkers* default settings (Data File S1).

### Differential Expression Analysis

Differential gene expression was conducted using the default limma-trend pipeline within each scRNA-seq batch to avoid batch effects (*51*). Subsequent gene set enrichment analyses based on all differentially expressed genes were conducted with the Gene Ontology, Reactome, and Hallmark databases using the *clusterProfiler* R package. Cell-cell interaction analysis was carried out using *CellChat* on all clusters present in individual day subsets.

### Tissue Culture of Organoids and Primary and Mouse Embryonic Fibroblasts

Mouse intestinal-derived organoids were cultured as previously described (*52, 53*). In brief, organoids were embedded in Matrigel (Corning, #356231) drops in 6 or 12 well-plate formats and grown in organoid media consisting of a 1:1 L-WNR conditioned medium and Advanced-DMEM/F-12 supplemented with HEPES, GlutaMax (Thermo Fisher, #35050061), N2 Supplement (Life Technologies, #17502048), Normocin (Invivogen, #ant-nr-1) and recombinant murine EGF (PeproTech, #315-09). Organoid plates were kept at 37°C and expanded every 3-4 days. Primary intestinal fibroblasts were cultured by plating single-cell solutions of dissociated lamina propria with DMEM/F-12 (Gibco #21041-025) supplemented with 10% FBS, 1X Pen/Strep, and 1μl/mL gentamycin (VWR #E737).

### In Vitro Interferon Treatment and In Vivo Interferon Blockade

*In vitro* interferon (IFN) treatment was performed by adding 500 U of mouse IFN-β (PBL Assay Science, #12401-1) to culture media of two-days grown MEFs for up to 6 hours. Cells were harvested for downstream RNA extraction and qPCR at indicated timepoints. For *in vivo* IFN-I blockade experiments, mice were treated with 500 mg/kg of anti-IFNAR-I blocking antibody (MAR1-5A3) at either 12 hours or 2 days after irradiation with FR or SR.

### RNA Extraction and Quantitative Real-Time PCR (qRT-PCR)

RNA was extracted using the Direct-zol RNA Microprep Kit (Zymo Research, #R2062). Briefly, single-cell pellets of dissociated intestinal tissue or organoids were resuspended in 500 μL or 250μL of TRIzol (Thermo Fisher, #15596018) respectively, and total RNA was isolated according to manufacturer’s instructions. NanoDrop spectrophotometry was used to quantify RNA concentration, and cDNA was subsequently generated from 500 ng of total RNA using Qscript cDNA synthesis kit (Quantabio, #95047-500). To analyze gene expression changes, qRT-PCR was performed using 3 ng of cDNA per reaction and in triplicates using Powerup SYBR master mix (Life Technologies, #A25778). SYBR primer sequences used in this study are annotated in (Data File S2). Gene expression was normalized to either peptidylprolyl isomerase A (*Ppia*) or glyceraldehyde-3-phosphate dehydrogenase (*Gapdh*).

### Immunofluorescence Staining in Intestinal Tissue Sections

For immunofluorescence of immune cell markers, the small intestine (jejunum) segments were harvested and preserved in optimal cutting temperature (OCT) medium. Jejunum segments were cut in 8-10-μm thick sections, coded, and stored at −80 °C. Slides were thawed at room temperature and subsequently fixed with 2% paraformaldehyde for 20 min. After three washes with Tris-Buffer saline (TBS), tissues were blocked with 8% bovine serum albumin (BSA) in TBS-T (0.025% Triton X-100) at room temperature for at least 1 h. The primary antibodies against MPO, F4/80, and CD3 were incubated overnight at 4 °C. After three washes with TBS-T, the secondary antibodies were added at 1:200 and incubated for 1h at room temperature in a humidified chamber. After 3 × 5 min rinses with TBS, tissues were stained with 1 μg/mL Hoechst (Invitrogen, H3570) for 30 min at room temperature and washed with TBS. Coverslips were mounted with antifade mounting medium.

For immunofluorescence of Olfm4, paraffin-embedded tissues were sectioned at 5µM thickness, placed on microscope slides, deparaffinized twice in Histo-Clear solution (Diamed, #NDIHS202) for 15min each, and rehydrated for 2mins in 100% ethanol (2x), 95%, and then 70%. Slides were then incubated for 15mins in 10mM Sodium Citrate (Sigma-Aldrich, #S4641-1KG) buffer supplemented with 0.05% Tween-20 (Millipore-Sigma, #P9416) at ∼95°C. Slides were cooled on ice for 10mins, washed for 5mins in 1X phosphate buffered saline (PBS) (Multicell, #311-425-CL), blocked at room temp for 1 hour in 1X PBS supplemented with 5% bovine serum albumin (BSA) (Sigma-Aldrich, #A7906) and 0.3% Triton X-100 (Sigma, #X100), and incubated overnight at 4°C with anti-Olfm4 (Cell Signaling Technology, # 39141S, 1:300) diluted in 1X PBS supplemented with 1% BSA and 0.3% Triton X-100. The next day, slides were washed thrice in 1X PBS for 5mins each and incubated for 3hrs at room temperature with DyLight650 (Invitrogen, SA5-10041, 1:500) and DAPI (Sigma-Aldrich, #D9542-1MG, 1:500) diluted in 1X PBS supplemented with 1% BSA and 0.3% Triton X-100. They were then washed three times in 1X PBS for 5mins each and mounted with Permount Medium (Fisher Scientific, # SP15-100). Images were acquired using a Nikon Ti2 inverted confocal microscope and analyzed using ImageJ software. Data is presented as percentage OLFM4+ crypts.

### RNA Scope in Intestinal Tissue Sections

smFISH (RNAScope; Advanced Cell Diagnostics) was performed on paraffin embedded sections according to the manufacturer’s recommendations. Tyramide signal amplification (TSA) fluorophores (Akoya Bioscience, TSA Fluorescein # NEL741001KT, TSA Cyanine 5# NEL745E001KT) were used for detection. The following probes were acquired from ACD: mm-Clu-C2(#427891-C2). Images were acquired at 20X magnification using Nikon Ti2 inverted confocal microscope.

### EdU Assay and detection

At 3.5 days post-irradiation, 200 mg of 5-ethynyl-20-deoxyuridine (EdU; Thermo Fisher Scientific, #C10337) dissolved in PBS were injected intraperitoneally 2 to 3 hours before euthanasia, as described previously (*34, 54*). EdU was detected according to the manufacturer’s instructions. Briefly, OCT embedded tissues were sectioned, fixed in 2% (w/v) PFA in PBS for 20 minutes at room temperature, washed twice with PBS, treated with 70% ethanol at -20°C, and washed with PBS three more times for 10 minutes each. Then sections were blocked in 8% bovine serum albumin (BSA) in PBS containing 0.5% Tween-20 and 0.1% Triton X-100 (PBS-TT) for at least 1 hour at room temperature in a humidified staining trough, followed by 5 minutes wash with PBS-TT. The Click-iT reaction mixture was then added to each slide and incubated at room temperature for 30 minutes. Each slide was washed in PBS-TT then counterstained with 100 microliters of Hoechst for 30 minutes, washed in PBS-TT, and mounted in Vectashield medium. Microscopy was performed on the Zeiss Observer Z1 (Jena, Germany).

### Quantification of OLFM4+ and Clu+ Cells

Quantification of Olfm4+ crypts/image was done manually using Image J software by counting the total number of positive crypts divided by the total number of visible crypts per image and presented as a percentage of Olfm4+ crypts. Quantification of Clu+ cells and Olfm4+ cells/crypt was done using QuPath software v4.0.3. In brief, to analyze the percentage of Clu+ cells/image, the entire image was first annotated. Next, DAPI+ cells were detected using the Positive Cell Detection plugin, which uses the "blue" channel to detect cells. After cell detection, a threshold of 5.0 was used for the “red” channel to detect Clu+ cells. Data is presented as a percentage of total Clu+ cells/image. To quantify the percentage of OLFM4+ cells/crypt, crypts containing OLFM4+ cells were first manually annotated for each image. Next, for each annotated crypt, DAPI+ cells were first detected, and then OLFM4+ cells were detected using a threshold of 400 for the DyL649 channel. Data is presented as a percentage of OLFM4+ cells/ crypt. A total of 25 images/mouse were quantified, and for each condition (NR, SR and FR), a total of 5 mice were analyzed.

### Quantification of Immune Cells

Quantification of staining for CD3 T cells, F4/80 (macrophages), and MPO (neutrophils) was performed manually using Image J software by counting the total number of positive cells divided by the total number of nuclei (DAPI) per 10x field. Data is presented as a percentage of positive cells. A total of 10 images/mouse were quantified and for each condition (NR, SR and FR) a total of 3-8 mice were analyzed (CD3, n=3 mice per condition; F4/80, n=7 mice for NR and n=8 mice for SR and FR; MPO, n=3 mice per condition).

### Quantification of Edu+ Cells

Quantification of average EdU+ cells per crypt was done manually using Image J software by counting the total number of positive EdU cells within each crypt. Data is presented as the average number of EdU+ cells/crypt per mouse. A total number of 100 crypts from 10 fields of view (10x magnification) per mouse were quantified. For each condition (SR and FR), a total of 5 mice were analyzed. In the case of the NR, 50 crypts in total (5 fields) were analyzed. In Figure 5D, one sample from the SR group was excluded due to poor tissue structure.

### Quantification of Intestinal Organoids

The number of live organoids were manually counted in images taken at days 0, 1, 3, and 5 post-irradiation. Data is presented as the proportion of surviving organoids, using the number of live organoids pre-treatment as a baseline.

### Statistical Analysis

Data are summarized using means, with SEM as error bars unless otherwise noted. Fisher exact testing was utilized to compare count data, while t-tests were used for pairwise comparisons of means. For comparisons involving three or more groups, ANOVA followed by post-hoc Tukey-HSD testing was used to determine significance among group means for normally distributed data. Otherwise, Kruskal-Wallis testing was used as the non-parametric equivalent of ANOVA. Two-factor ANOVA followed by post-hoc estimated marginal means was used to analyze EdU data obtained from the anti-IFNAR experiment. Linear mixed effects models were used to analyze post-irradiation weight and organoid survival data over time and across treatment modalities. The Gehan-Breslow-Wilcoxon method was used to compare survival data, while the Cox proportional-hazards model was used to determine significant co-variates impacting survival.

### Linear Regression Analysis

Scaled single-cell expression data for control and day 2 irradiated samples were used to assess the independent effects of IFN-I signaling on pathways scores for Hallmark gene sets and select REACTOME pathways. For each cell type from the expression data, the *Seurat* module scores for the Hallmark IFNA and IFNG gene sets together with the treatment group (control, FR, SR) were used in a full linear regression model, treating the module scores for each Hallmark or REACTOME gene set of interest as the response variable. For each cell type, this resulted in one model for each pathway of interest. Each model was then filtered by significance of the SR treatment group level (p < 0.01), the FR treatment group level (p < 0.10), and minimum FR effect size coefficient (0.30), using the control treatment group as the reference level for the model. This resulted in cell types and pathways for which IFN-I pathway scores exhibited differential effects between FR and SR, controlling for the effect of IFNG pathway scores. Marginal effects for variables in the model were determined using the *sjPlot* R package.

## List of Supplementary Materials

Supplementary Figure 1

Supplementary Figure 2

Supplementary Figure 3

Supplementary Figure 4

Supplementary Figure 5

Supplementary Figure 6

Data File S1: List of genes differentially expressed across all cell clusters identified in the scRNA-seq analysis.

Data File S2: RT-qPCR primers used in this study.

## Acknowledgments

The irradiations were performed by the Cell and Animal Radiation Core Facility (RRID:SCR_022377) at the University of Pennsylvania Perelman School of Medicine.

## Funding

This work was supported by NIH grant 5P01CA257904-02 to CK and AJM and the Mark Foundation for Cancer Research to AJM. This work was partially supported by a grant to I.I.V. by the National Center for Advancing Translational Sciences of the National Institutes of Health under award number UL1TR001878 and the Institute for Translational Medicine and Therapeutics’ (ITMAT) Transdisciplinary Program in Translational Medicine and Therapeutics. The content is solely the responsibility of the authors and does not necessarily represent the official views of the NIH.

### Author contributions

Conceptualization: CK, AJM

Methodology: TLL, CM, IIV, KK, LL, CJF, KN, MP, AV, PC, SB, JLW

Investigation: TLL, CM, IIV, KK, LL, CJF, MMK, BY, KN, SAOM, AV, PC, SB, DG

Visualization: TLL, CM, IIV, AJM Funding acquisition: CK, AJM

Supervision: CK, AJM

Writing – original draft: TLL, CM

Writing – review & editing: TLL, CM, IIV, KK, LL, CJF, MMK, NL, BY, KN, MP, SAOM, AV, PC, SB, DG, CJL, JLW, CK, AJM

## Data and materials availability

All data associated with this study are present in the paper or the Supplementary Materials. The datasets for single-cell sequencing generated and analyzed during the current study are available in the Gene Expression Omnibus (GEO) repository under accession number ***.

**Figure S1.**
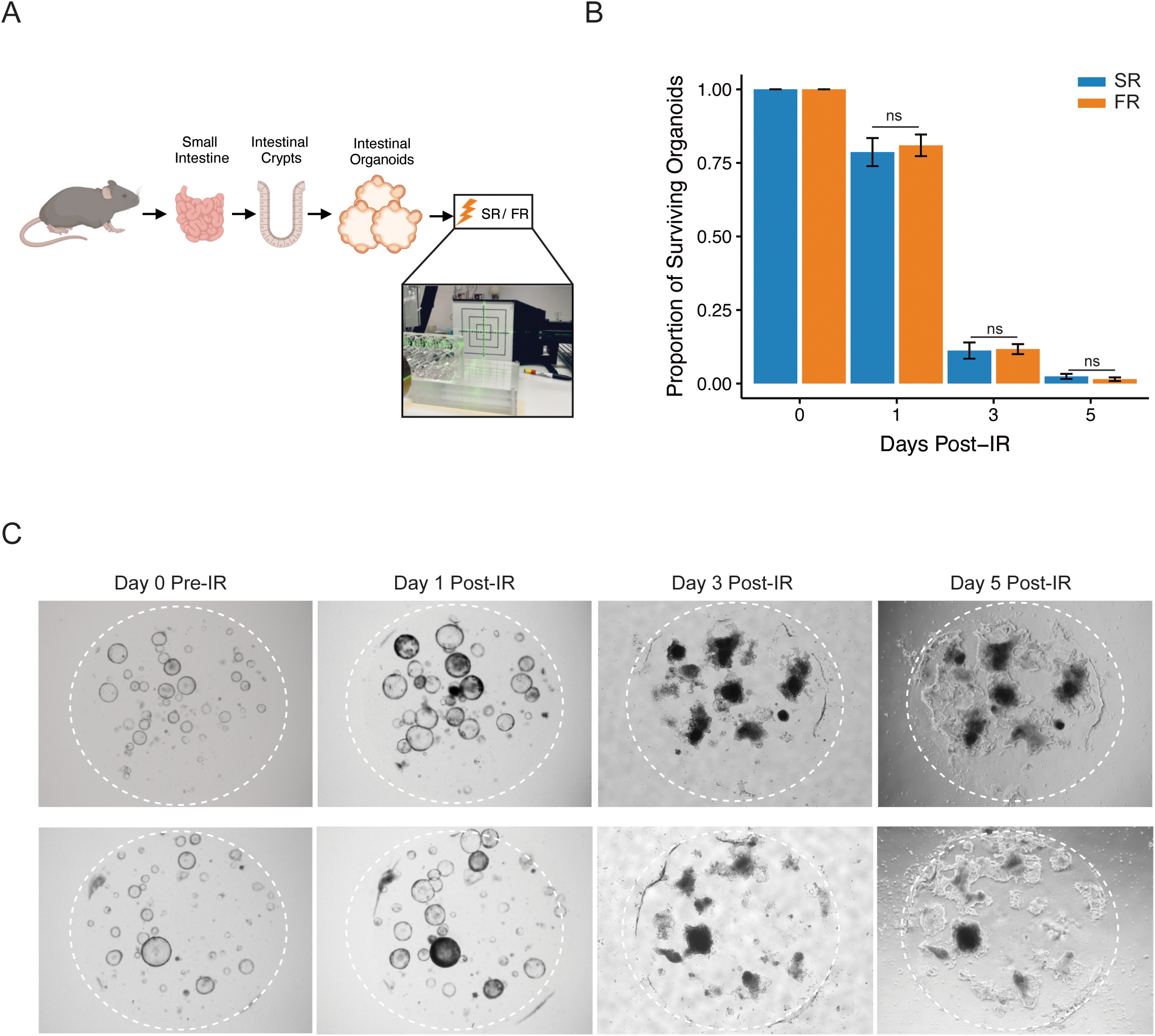
The FLASH effect is dependent on the tissue microenvironment. **(A)** Schematic representation of intestinal organoid isolation and *ex vivo* irradiation process. **(B)** Average proportion of surviving organoids at specified time points after single-dose of SR or FR. The bars represent the proportion of surviving organoids, and error bars indicate the standard deviation. (n=9 images were quantified for each time point and condition). **(C)** Representative image of *in vitro* intestinal mouse organoids at indicated time points after SR or FR.

**Figure S2.**
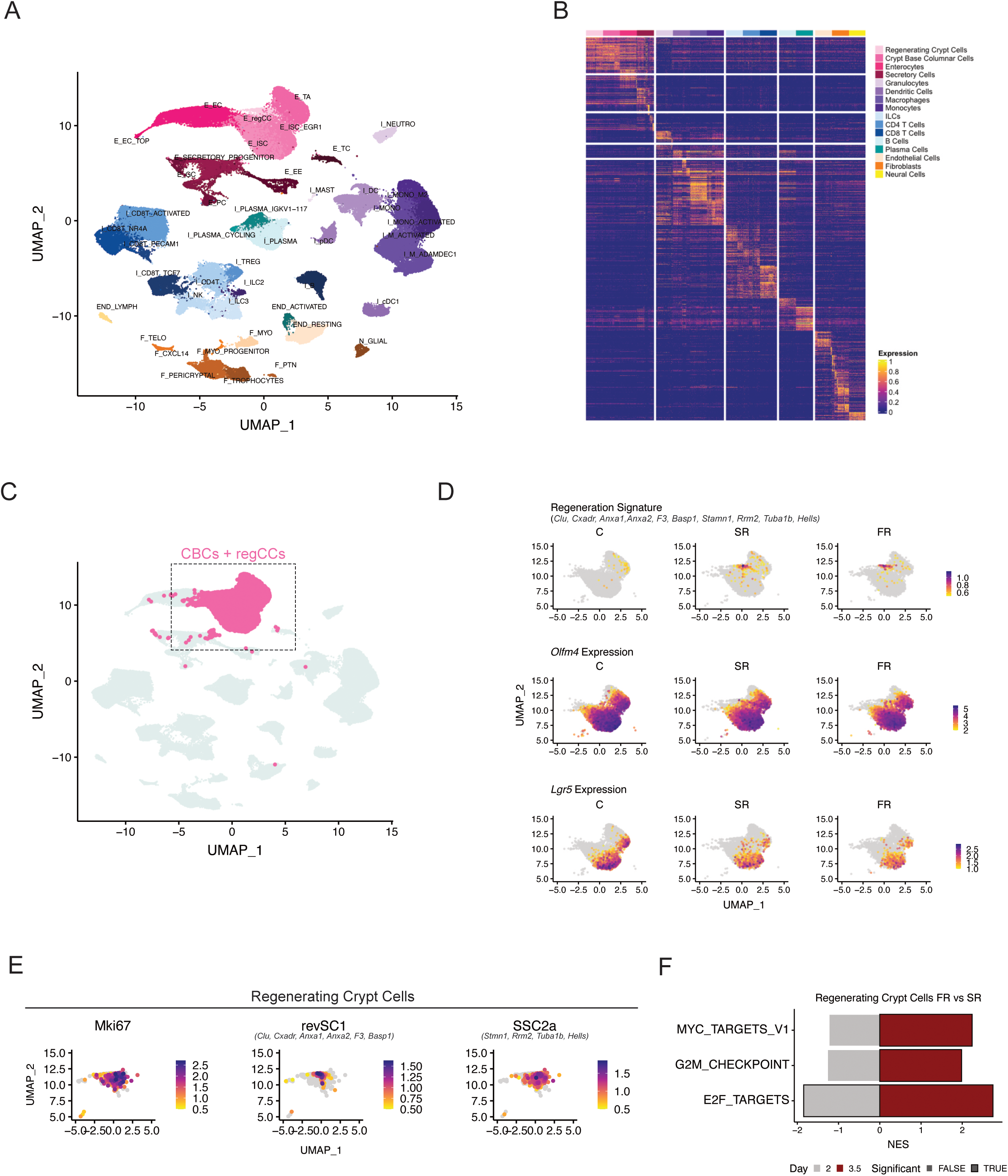
Time course scRNA-sequencing captures intestinal regeneration and recapitulates the FLASH effect. **(A)** UMAP plot of epithelial and lamina propria cell clusters at all time points color-coded by cell sub-type. **(B)** Heatmap showing expression of top differentially expressed genes across all cell types annotated in the single-cell RNA sequencing (scRNA-seq) data (Supplementary Data 1). **(C)** Crypt base columnar (CBC) cells and regenerating crypt cells (regCCs) in the UMAP plot. **(D)** Expression of regenerative gene signature (Supplementary Data 2) and stem cell markers Lgr5 and Olfm4 across CBCs and regCCs clusters in non-irradiated (NR), SR, and FR samples. **(E)** Expression of proliferative gene mki67 or revSC1 and ssc2a1 gene signatures (Supplementary Data 2) in the regCCs in NR, SR, and FR samples. **(F)** Gene set enrichment analysis of proliferation-related gene sets within the regCCs from mice treated with FR compared to SR at day 3.5 post-RT.

**Figure S3.**
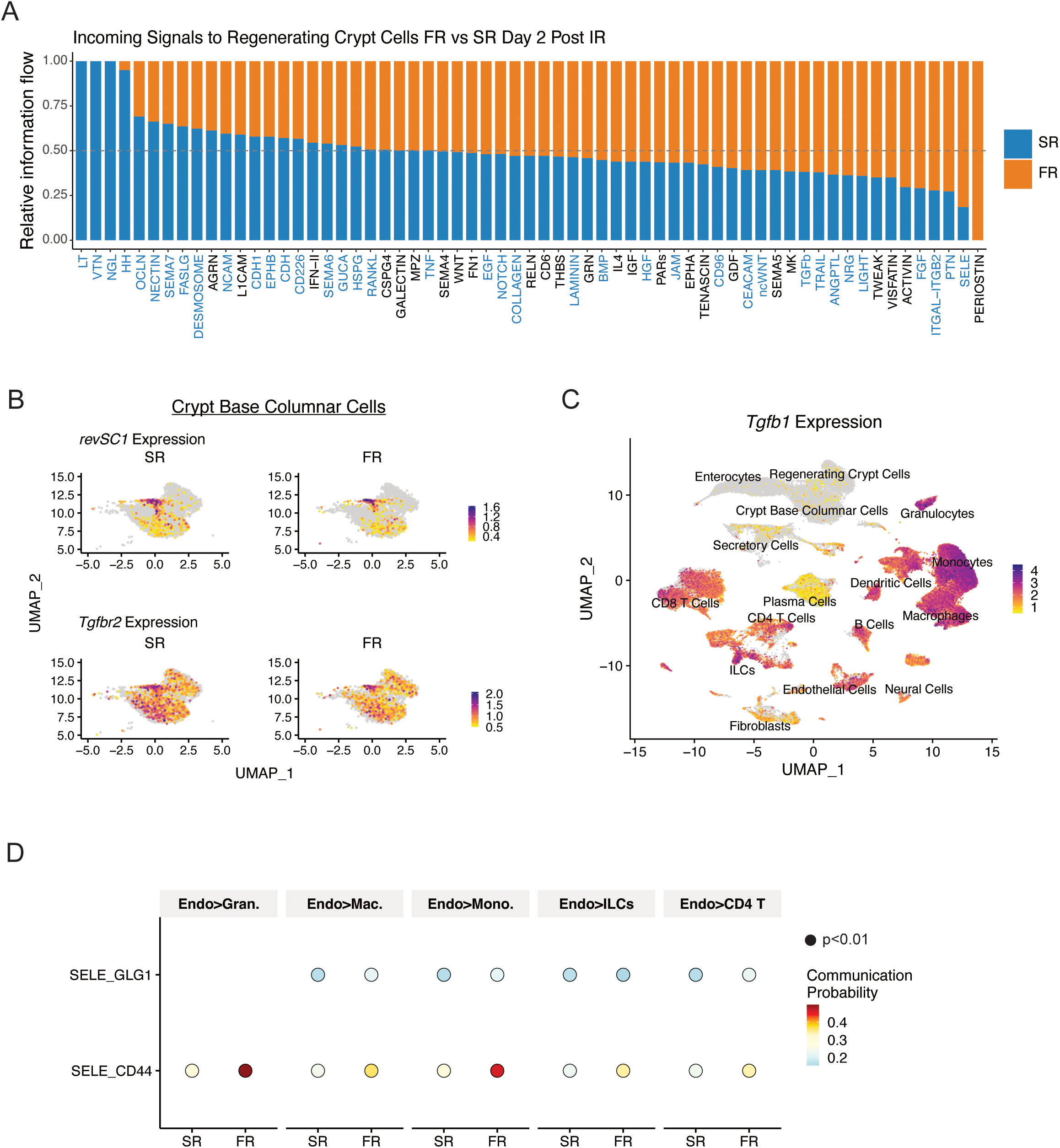
FLASH RT is associated with higher immune cell infiltration and TGFb signaling in the regCCs. **(A)** Stacked bar plot comparing the incoming signals to regCCs between FR and SR two days after RT. Signaling pathways on the x-axis that are highlighted represent significantly different pathways between FR and SR. **(B)** UMAP showing the average expression of the revSCs gene signature and the *Tgfbr2* gene in the crypt base columnar cells in SR and FR samples from all time points. **(C)** UMAP showing the expression of Tgfb1 ligand in all cell types and time points in SR and FR samples. **(D)** Bubble chart showing E-selectin ligand-receptor pair signaling from endothelial cells to immune cells in SR and FR samples, calculated by CellChat. Dot color reflects communication probabilities, and dot size represents computed p-values.

**Figure S4.**
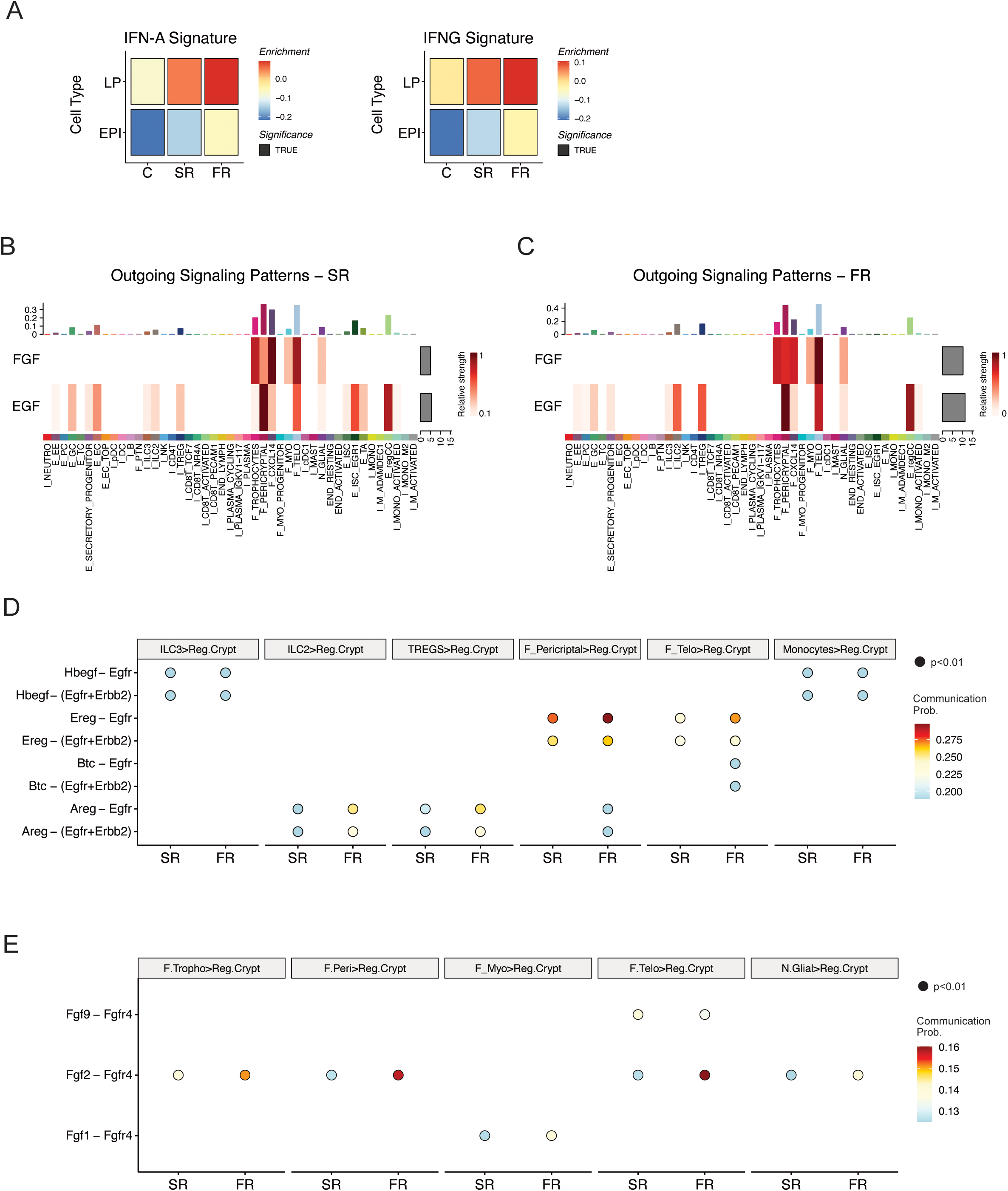
FLASH RT is associated with higher IFN signaling and growth factor production in the stromal cells from irradiated intestines. **(A)** Relative expression of IFN alpha and gamma gene sets in FR and SR samples two days post-RT, faceted by epithelial and lamina propria cell compartments. **(B, C)** Heatmaps of EGF and FGF signals contributing to outgoing signaling across all cell types in SR (left) or FR (right). The gray bars indicate the sum of the signaling strength of each signaling pathway across all cell types. **(D)** Bubble chart showing EGF ligand-receptor pair signaling from immune cells and fibroblasts to regCCs calculated by CellChat in SR and FR. Dot color reflects communication probabilities, and dot size represents computed p-values. **(E)** Bubble chart showing FGF ligand-receptor pair signaling fibroblasts to regCCs calculated by CellChat in SR and FR. Dot color reflects communication probabilities, and dot size represents computed p-values

**Figure S5.**
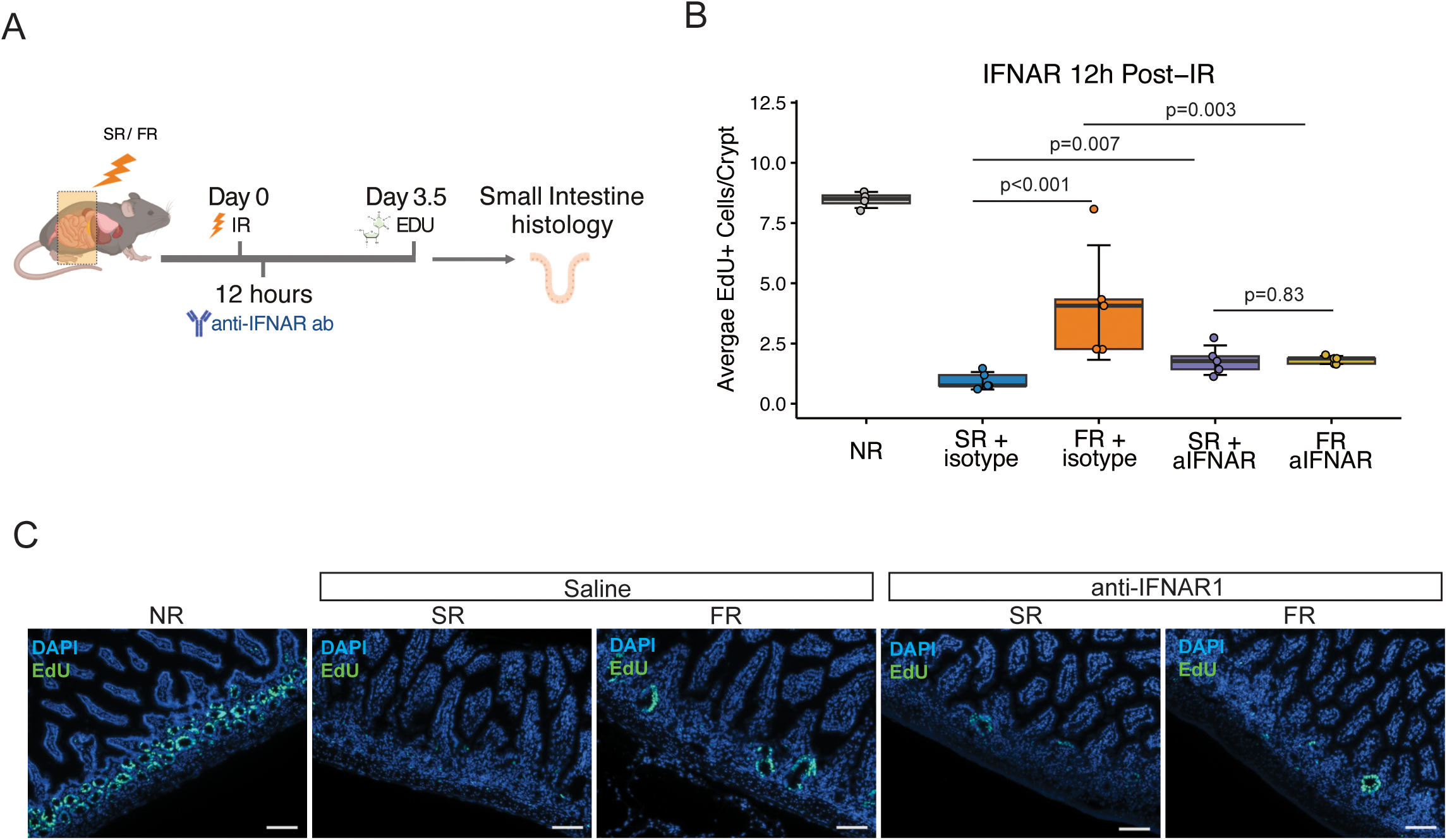
FLASH effect is abolished with early administration of IFNAR1 blockade. **(A)** Schematic representation of whole abdomen RT and follow-up assays. Anti-IFNAR antibody was administered 12h post-RT. **(B)** Average number of EdU+ cells per crypt in the intestines of mice treated with no irradiation (NR), SR, or FR at day 3.5 post-treatment with or without IFNAR1 blocking antibody given 12h post-treatment. Boxplot represents interquartile range (IQR) with median indicated. Whiskers represent highest and lowest values within 1.5 times the IQR. Each dot is the average of 10 tissue areas quantified per mouse (n=5 biologically independent samples per group). Statistical significance was calculated using two-factor ANOVA testing followed by estimated marginal means. **(C)** Representative immunofluorescent images from isolated small intestines stained for EdU from NR or at Day 3.5 following SR or FR with or without IFNAR1 blocking antibody treatment. Magnification, ×10; Scale bars, 100 μm.

